# Impacts of climate change on the movement ecology of an imperfect homeotherm

**DOI:** 10.1101/2025.02.24.639910

**Authors:** Michael J. Noonan, Jesse M. Alston, Aline Giroux, Arnaud L.J. Desbiez

## Abstract

Rapid, human-induced climate change has posed significant challenges to wildlife. One key strategy animals use to cope with environmental temperature fluctuations is behavioral thermoregulation. Understanding how climate change is expected to influence animal behavior is crucial for assessing its impact on species survival and informing effective conservation efforts. Giant anteaters have been found to exhibit conspicuous behavioral responses to temperature changes. Despite their broad thermal neutral zone (15 – 36°C), climate projections indicate that this vulnerable mammal is increasingly likely to experience heat stress. We used GPS tracking and continuous-time analyses to investigate how environmental temperature influences the movement ecology of giant anteaters. We integrated our findings with climate change projections to link giant anteater’s responses to present weather conditions with those expected under future climate scenarios. Giant anteaters’ movement speed exhibited a negative quadratic response to temperature, peaking at 23.7°C. 95% of their movement occurred between 15.0 – 32.3°C, which aligns with their thermal neutral zone. The increasing temperature led giant anteaters to increase selection for native forests, but had no effect on selection for exotic tree plantations. This shows the importance of native forests as these thermal shelters help to mitigate the negative consequences of high temperatures on anteater’s movement. However, the warmer temperatures predicted for Brazil throughout the rest of the 21st century indicate that giant anteaters may experience a reduction of up to 84% in their movement speed. This would hinder the acquisition of sufficient energy resources and threaten the species’ persistence. We emphasize the need for conservation efforts that account for the impacts of climate change on species survival and stress the importance of preserving forests as essential refuges that help wildlife to cope with rising temperatures.

## Introduction

It is now widely recognised that human-induced rapid environmental change (Sih *et al*., 2011) has triggered the Earth’s sixth major episode of mass extinction (Barnosky *et al*., 2011, Ceballos *et al*., 2015, 2017). Although the ongoing biodiversity crisis has been caused primarily by habitat loss (Venter *et al*., 2006, Woo-Durand *et al*., 2020), it is being compounded by rapid, human-induced climate change (Mantyka-Pringle *et al*., 2015, Woo-Durand *et al*., 2020), which is accelerating extinction risks globally (Urban, 2015, Wiens, 2016). Consensus is growing among ecologists that behavioral responses will play a major role in influencing species’ susceptibility to climate change (Beever *et al*., 2017, Huey *et al*., 2012, Kearney *et al*., 2009, Moritz & Agudo, 2013, Sunday *et al*., 2014). Nevertheless, we still know little about species’ behavioural responses to thermal stress and other acute weather events and the consequences of these responses for fitness (Buchholz *et al*., 2019), particularly in large, free-ranging mammals.

From the research on this topic that has been performed, it is clear that wildlife rely on behavioral plasticity to balance energetic intakes and expenditures during shifting weather conditions. For instance, many species reduce their activity (Alston *et al*., 2022, Giroux *et al*., 2021a, Noonan *et al*., 2014, Sheppard *et al*., 2021), modify their circadian rhythms (Camilo-Alves & Mourão, 2006, Giroux *et al*., 2023, Levy *et al*., 2019), and/or adjust their use of thermal refugia across the landscape (Alston *et al*., 2020, Long *et al*., 2014, Sarmento *et al*., 2019) during periods of adverse weather to save energy on thermoregulation. During historically normal climatic cycles, this behavioral plasticity likely provided individuals with the capacity to flexibly minimize the energetic costs associated with thermoregulation. Over long periods of atypical weather, however, such as those induced by anthropogenic climate change, these once-useful behaviors risk becoming unsustainable if they do not result in sufficient energetic gains (e.g., Cunningham *et al*., 2020, Haase *et al*., 2020, Noonan *et al*., 2018). By studying how individuals adjust their behavior and balance energetic trade-offs in response to changes in weather, we can therefore make predictions about how wildlife might respond to future climate change and better support proactive conservation efforts. Developing a full understanding of the role of behavioural plasticity in managing thermal stress will require studies on both thermal specialists and generalists. Giant anteaters are (*Myrmecophaga tridactyla*) expected to be comparatively less sensitive to heat stress than other comparably-sized mammals McNab (1984, 1986), making them a useful model for evaluating the impact(s) of climate change on mammalian movement behaviour.

The impacts of ongoing and future climate change on giant anteater behaviour are of particular conservation concern. Giant anteater populations have suffered severe reductions in recent years (including local and regional extirpations) and are currently classified by the International Union for Conservation of Nature (IUCN) as Vulnerable (Miranda *et al*., 2014). While habitat loss and road-induced mortality are currently recognised as the primary threats to the species’ survival, their capacity to cope with future climate change will play an important role in dictating population trends. Distributed throughout Central and South America (Gaudin *et al*., 2018), the species originated during the late Paleocene (O’Leary *et al*., 2013) and has persisted through numerous periods of extreme climatic change — such as the global cooling and aridisation that occurred during the Eocene-Oligocene transition (Prothero, 1994). Giant anteaters exhibit a metabolic rate that is 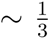 of that expected by Kleiber’s rule (Kleiber, 1932, McNab, 1984), resulting in low body heat production and, consequently, low body temperatures (McNab, 1984). This likely makes them less sensitive to high ambient temperatures compared to other mammals of similar body mass. Additionally, giant anteaters have a much lower thermal conductance than other similarly sized tropical mammals (McNab, 1984), allowing them to remain thermally neutral over a much broader temperature range (∼15 – 36°C) than other ant- and termite-eating specialists (McNab, 1984). Despite this, a growing body of literature demonstrates that giant anteaters currently exhibit conspicuous behavioral responses to cope with weather changes (Camilo-Alves & Mourão, 2006, Giroux *et al*., 2023, 2021a, Mourão & Medri, 2007). Furthermore, the warmer temperatures predicted under probable climate change scenarios (Masson-Delmotte *et al*., 2021) suggest that giant anteaters are increasingly likely to experience conditions that induce heat stress (≥36°C). Yet, individuals will face a trade-off between investing in behavioral thermoregulation versus other essential activities such as feeding or mating (*sensu* Cunningham *et al*., 2020). It is thus important to predict how giant anteaters’ behaviour is expected to respond to human-induced rapid climate change and understand how these responses might affect individual fitness and population viability.

Using a two-step approach, we aimed to predict the impacts of future climate changes on giant anteater movement ecology. We first used GPS location data and continuous-time analyses (Alston *et al*., 2023, Noonan *et al*., 2019a) to estimate the influence of environmental temperature on anteaters’ i) instantaneous movement speeds; and ii) habitat selection. Because of giant anteaters’ low thermal conductance (McNab, 1984), we expected that individuals would decrease their movement speed and increase their selection for thermal refugia during both hot and cold spells, as compared to thermally normative periods (Camilo-Alves & Mourão, 2006, Giroux *et al*., 2023, 2021a, Mourão & Medri, 2007). We then paired these findings with predictive models based on the Intergovernmental Panel on Climate Change’s (IPCC) Shared Socioeconomic Pathways (SSP) climate change scenarios (Riahi *et al*., 2017) to link the mechanistic responses to current weather to conditions expected under future climate change. We anticipated that the warmer temperatures predicted under probable climate change (Masson-Delmotte *et al*., 2021) would lead to giant anteaters reducing their movement speed and increasing their selection for thermal refugia. These analyses allow for in-depth explorations of how predicted climate change may drive adaptive changes in the movement ecology, space use, and energetics of giant anteaters.

## Methods

### Study area

The study was conducted at three sites in the state of Mato Grosso do Sul (MS), in the Cerrado biome (savannah) of Brazil (Fig. 1). The climate throughout MS is wet from October to March and dry from April to September (Köppen’s Aw), with historically mild year-round temperatures (range 21-32°C), though temperatures can exceed 40°C in the summers and drop to 0°C in the winters (Alvares *et al*., 2013). Land use is dominated by pasture, with sparse remnants of native forest and savanna, and some areas of eucalyptus plantation. Streams bordered with native riparian vegetation are common throughout the study area.

**Figure 1:**
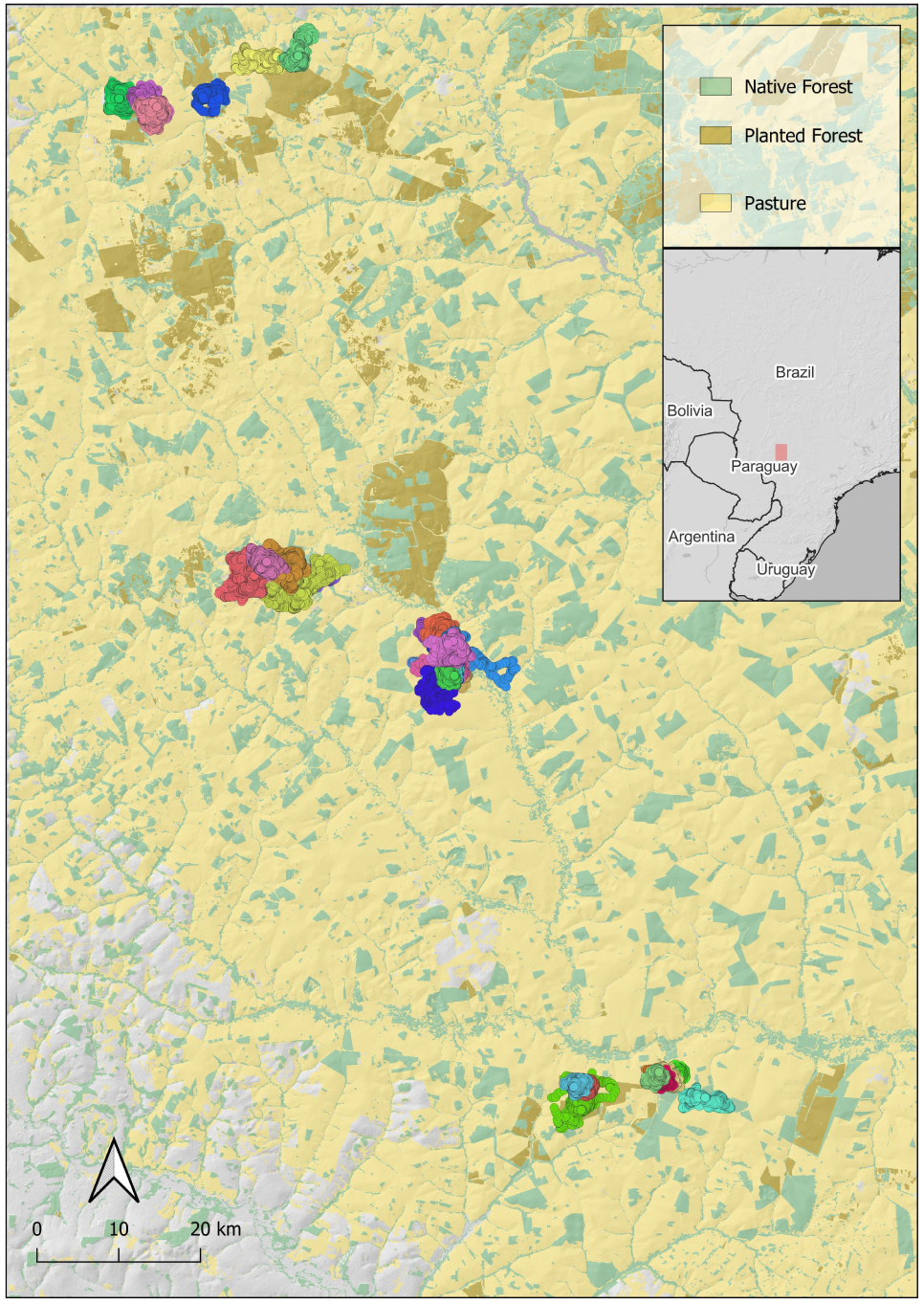
Map showing the GPS tracking data for the monitored animals in Mato Grosso do Sul, Brazil. The different colours correspond to location data from different individuals. The native forest, planted forest, and pasture MapBiomas land cover classifications for the region are also shown.

### Giant Anteater Tracking Data Collection

Free-ranging giant anteaters were captured between 2017 and 2018 within the three study sites. All animals were captured at distances of >15 km from urban areas, and equipped with GPS tracking collars. The capture team was always comprised of two veterinarians and a biologist, minimum. To capture giant anteaters, we searched open areas for foraging individuals, during colder months (May-August) when individuals are known to exhibit greater diurnal activity (Camilo-Alves & Mourão, 2006). When an adult was spotted, it was approached on foot by two members of the team and captured using two long-handled dip-nets (handle 1.5m; hoop 0.7m diameter) to restrain it. A veterinarian then applied an intramuscular injection of butorphanol tartrate (0.1mg/kg), detomidine hydrochloride (0.1mg/kg) and midazolam hydrochloride (0.2mg/kg) into the animal’s hind limb. After anaesthetic induction, the front claws were first wrapped and completely immobilized using tape. Physical exams were then performed to evaluate animal health, including measuring weight, reproductive status (via stomach palpation), general appearance, hydration status, mucous membrane colour, respiratory auscultation, and presence of scars or wounds. Adult individuals assessed by veterinarians to be in good health were fitted with a GPS harness (TGW-4570-4 Iridium GPS) and VHF transmitter (MOD 400; Telonics, Mesa, Arizona). For anaesthetic reversal procedures, all individuals received a combination of three antagonists: naloxone hydrochloride (0.02mg/kg), yohimbine hydrochloride (0.125mg/kg), and Flumazenil (0.01mg/kg). After the procedure, the animal was maintained in a wooden ventilated crate until complete recovery and was then released at its capture location. Full details on the capturing and handling protocols are detailed in Kluyber *et al*. (2021). Collared giant anteaters were recaptured approximately one year afterward for harness removal and data download, but each animal was visually inspected at a distance through binoculars at least once every two weeks for a general health check.

### Data Analysis

#### Movement data pre-processing

Before analysis, we performed a data cleaning process in order to calibrate the GPS measurement error and remove any outlying data points. This was done by using the methods implemented in the R package ctmm (version 0.5.3; Calabrese *et al*., 2016). For each location estimate, the GPS trackers recorded a unitless Horizontal Dilution of Precision (HDOP) value which is a measure of the accuracy of each positional fix. We converted the HDOP values into calibrated error circles by estimating an equivalent range error from 6,948 calibration data points where a tag had been left in a fixed location (Fleming *et al*., 2021). For each individual animal, we then removed outliers based on error-informed distance from the median location, and the minimum speed required to explain each location’s displacement. For further details on the pre-processing see Appendix S2 in Noonan *et al*. (2022).

#### Movement Analyses

For each individual anteater, we fitted a continuous-time movement model to the location data and conditioned our movement analyses upon the best model for each individual. We used variogram analysis (Fleming *et al*., 2014) to ensure animals were range-resident, then fitted and selected an autocorrelated movement model that best described the animal’s movements using perturbative Hybrid Residual Maximum Likelihood (phREML; Fleming *et al*., 2019) and Akaike’s Information Criterion corrected for small sample sizes (AICc).

From each of the best-fit models we then estimated the instantaneous movement speeds (in m/s) at each sampled time-point, using continuous-time speed and distance estimation (Noonan *et al*., 2019b). This method uses a simulation-based approach to sample from the distribution of possible trajectories that are consistent with the data and a fitted continuous-time movement model, to generate an estimate of the mean speed ± confidence intervals. This approach corrects for GPS measurement error and is insensitive to the sampling schedule, enabling robust comparisons.

To estimate 3^rd^ order habitat selection (*sensu* Johnson, 1980), we used weighted resource selection functions (Alston *et al*., 2023). This method borrows likelihood weights from home range estimates generated using weighted Autocorrelated Kernel Density Estimation (AKDE) (Fleming *et al*., 2018) and applies them to individual locations, which enables more robust estimates of uncertainty around selection parameters for individual animals than other currently available habitat selection analyses. We sampled availability for each animal using an integrated model, fitting a Gaussian model of availability simultaneously with estimation of resource selection parameters. The probability density function of the model can be represented as

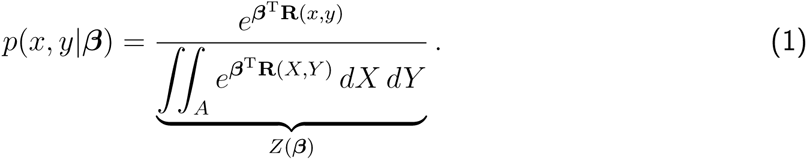

where **R**(*x, y*) represents spatially explicit covariates or resources, *β* is their corresponding regression parameters, *A* is the area over which availability is sampled, and *Z*(*β*) is a normalization constant. In our model,

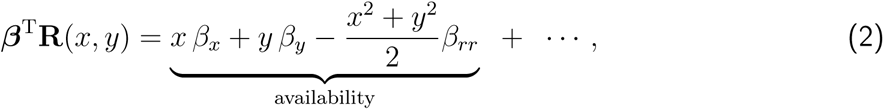

where the availability terms highlighted in Eq. (2) set the area in which availability is sampled (i.e., the sampling window), followed by additional resource selection parameters. This method of sampling availability in RSFs is conceptually similar to availability sampling in integrated step-selection analysis (Avgar *et al*., 2016).

We included in our model two environmental covariates that may represent features of the landscape used by anteaters as thermal cover: native forest and eucalyptus plantations. To examine how temperature influences use of these environmental covariates, we also included interaction terms between temperature (ECMWF ERA5-Land hourly temperature, 2 m above surface; Muñoz-Sabater *et al*., 2021) and the two covariates of interest. Native forest and plantation were determined using collection 7, 30m resolution MapBiomas land cover data (Souza *et al*., 2020), with native, Cerrado forest defined as ‘*vegetation types with a predominance of tree species with continuous canopy formation*’, and plantations as ‘*tree species planted for commercial purposes*’. We standardized temperature across the entire data set of anteater locations to aid integration of the resource selection function. We generated two RSFs for each individual— one RSF parameterized using daytime locations, and a second parameterized using nighttime locations—and used AICc to select the best set of parameters from the global model. We then generated population-level estimates of habitat selection for day and night from the parameter estimates of individual resource selection functions using log-normal meta-analysis.

For all movement analyses, we used the ctmm R package (v1.2.1; Calabrese *et al*., 2016) in the R statistical software environment (v4.3.1; R Core Team, 2020).

#### Climate Change Projections

We used the outputs of our empirical analyses to inform predictions of how future climate change could affect giant anteater movement behavior. Because our primary aim was to assess the species’ vulnerability to future climate change, we kept our projections deliberately coarse and focused only on predicting mean monthly movement rates. Using the World Bank’s Climate Knowledge Portal web interface, we obtained projected mean monthly temperature data for the state of Mato Grosso do Sul over the coming century. Temperature data were projected based on the global climate model compilations of the Coupled Model Inter-comparison Projects (CMIPs), overseen by the World Climate Research Program. Projections were obtained for five Shared Socioeconomic Pathways (SSP) scenarios (SSP1-1.9, SSP1-2.6, SSP2-4.5, SSP3-7.0, and SSP5-8.5) that support the IPCC’s Sixth Assessment Report (Masson-Delmotte *et al*., 2021). For each scenario, we estimated the movement speeds based on the mean annual temperature predicted to occur between 2014–2100. To obtain these estimates we generated a sample of 10,000 bootstrapped estimates of movement speeds for each of the predicted mean annual temperatures by re-sampling from the empirical instantaneous movement speed data. We then estimated the mean movement rate, in km/day, from this sample. As an example of this process, under SSP1-1.9, the mean annual temperature for 2014 was predicted to be 25.9°C. We therefore re-sampled 10,000 estimates of the movement speeds that occurred at 25.9°C during the study period. This resulted in a projected mean movement speed of 3.28 km/day (95% CI: 3.16 – 3.40 km/day). We note that although the general trends can be considered to be robust, both our empirical estimates and the climate predictions are subject to modeling error, and exact values should be treated with caution.

## Results

We deployed collars on 43 individuals. Two of the collared animals had insufficient data due to collar malfunctions, and three of the individuals dispersed over the study period, and were therefore excluded from our analyses as we were interested primarily in understanding typical behaviour from range-resident individuals. We therefore present results for 38 range-resident giant anteaters. Tags operated for a median of 11.2 months across all tagged giant anteaters and took GPS fixes at 20-min intervals. The final GPS dataset comprised 847,683 GPS fixes collected over 12,761 individual-days (Fig. S1).

### Temperature and movement rates

Over the duration of the study period we monitored giant anteater movement between temperatures of 1.9 – 39.9 °C. Estimated giant anteater movement and activity showed clear relationships with temperature. Movement speed ranged between 0 and 0.86 m/s, with the peak of their movement occurring at 23.7°C and 95% of their movement occurring between 15.0 – 32.3 °C (Fig. 2A). Indeed, the probability of giant anteaters being active or not was best described by a quadratic relationship with temperature (Table 1, Fig. 2B). This relatively simple model was able to predict activity patterns with an accuracy of 63%, demonstrating the importance of temperature as a driver of giant anteater behaviour.

**Figure 2:**
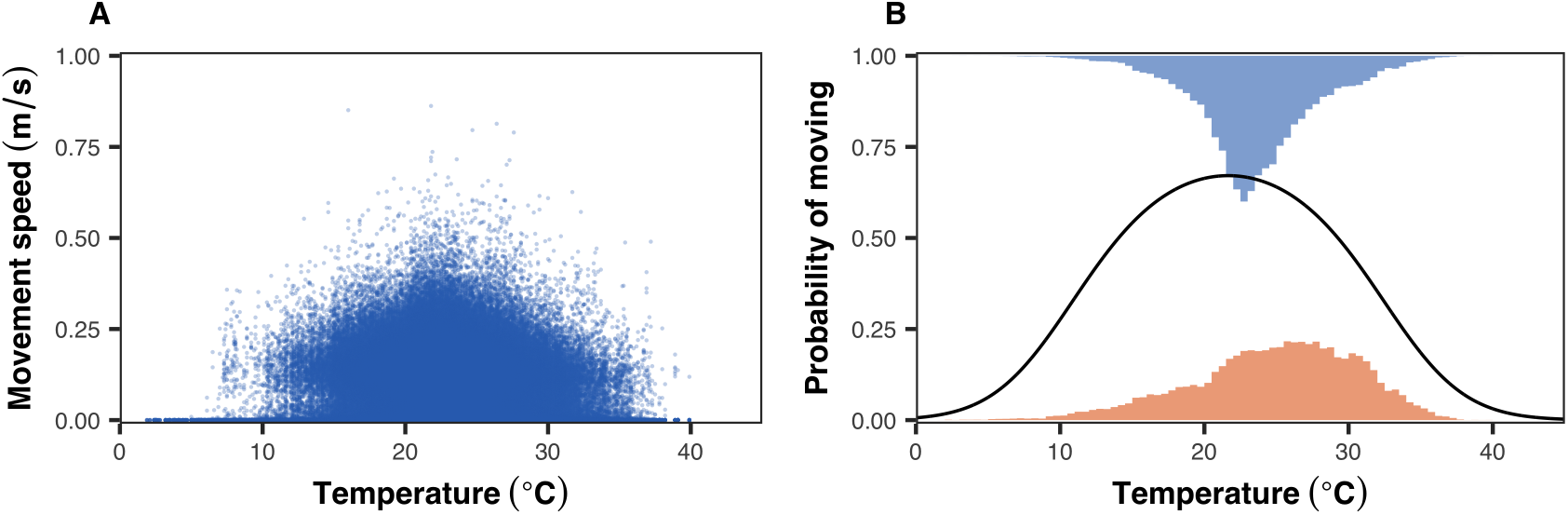
The relationship between anteater movement and temperature. The scatterplot in (**A**) depicts instantaneous movement speed estimates as a function of the air temperature at the time of the estimate (*n*= 758,045). Panel (**B**) shows the results of a binomial regression model predicting whether giant anteaters were active or not as a function of air temperature. The histograms show the distribution of temperatures over which individuals were classified as being active (blue), or inactive (orange).

**Table 1:**
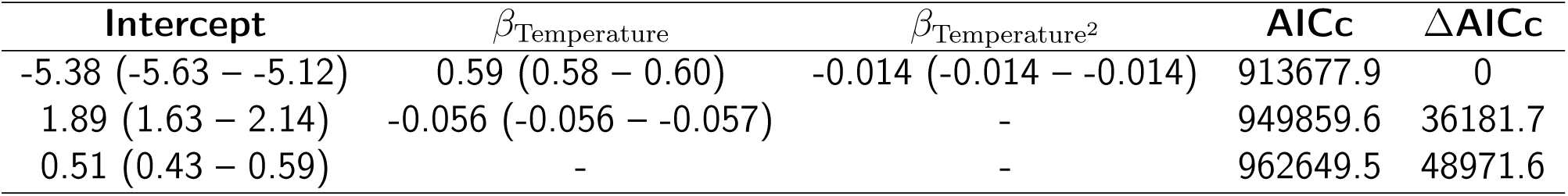
Model selection results for determining the extent to which giant anteater activity is governed by air temperature. The best fit model was identified by Akaike’s Information Criterion corrected for small sample sizes (AICc). All models included a random intercept around individual giant anteaters. Parameter values are shown on the logit scale.

| Intercept | $\beta_{\text{Temperature}}$ | $\beta_{\text{Temperature}^2}$ | AICc | $\Delta\text{AICc}$ |
| --- | --- | --- | --- | --- |
| -5.38 (-5.63 – -5.12) | 0.59 (0.58 – 0.60) | -0.014 (-0.014 – -0.014) | 913677.9 | 0 |
| 1.89 (1.63 – 2.14) | -0.056 (-0.056 – -0.057) | - | 949859.6 | 36181.7 |
| 0.51 (0.43 – 0.59) | - | - | 962649.5 | 48971.6 |

### Temperature and habitat selection

We found empirical support for selection of native forest and the interaction between temperature and native forest in our population-level resource selection functions for both day and night (Table 2), indicating that anteaters selected native forest and increased selection of native forest as temperatures increased. We detected empirical support for selection of plantation during the day but not night, and we did not detect support for an interaction between temperature and plantation during either day or night.

**Table 2:** Results of population-level resource selection functions. Each cell contains the model coefficient for a covariate, followed by a 95% confidence interval in parentheses. The cells for interactions between plantation and temperature are empty because this interaction was never included in the most-supported model for any individual.

| Time of Day | Covariate | Coefficient | Interaction w/ Temperature |
| --- | --- | --- | --- |
| Day | Native Forest | 1.12 (0.62, 1.62) | 0.44 (0.31, 0.58) |
|  | Plantation | 1.49 (0.43, 2.54) | — |
| Night | Native Forest | 0.60 (0.14, 1.06) | 0.80 (0.60, 1.00) |
|  | Plantation | -0.02 (-1.01, 0.97) | — |

### Climate change projections

Predicted climate change through the rest of the twenty-first century appears likely to result in substantial reductions in giant anteater movement. The modestly warmer conditions predicted by the end of the century under the lower emissions SSP1-1.9 (mean annual temperature: 26.2 °C) and SSP1-2.6 scenarios (mean annual temperature: 26.7 °C) could cause net reductions in movement of 13.2% (95% CIs 9.9 – 16.5%) and 25.4% (95% CIs 22.2 – 28.5%) respectively (Fig. 3). The temperatures predicted under the higher emissions scenarios may drive dramatic reductions in activity. The SSP3-7.0 scenario (mean annual temperature: 29.7 °C), might bring about a 66.9% reduction in movement (95% CIs 64.7 – 69.0%), while giant anteater movement under the SSP5-8.5 scenario (mean annual temperature: 31.0 °C), risk being as much as 84.0% lower than present day rates of movement (95% CIs 82.4 – 85.5%).

**Figure 3:**
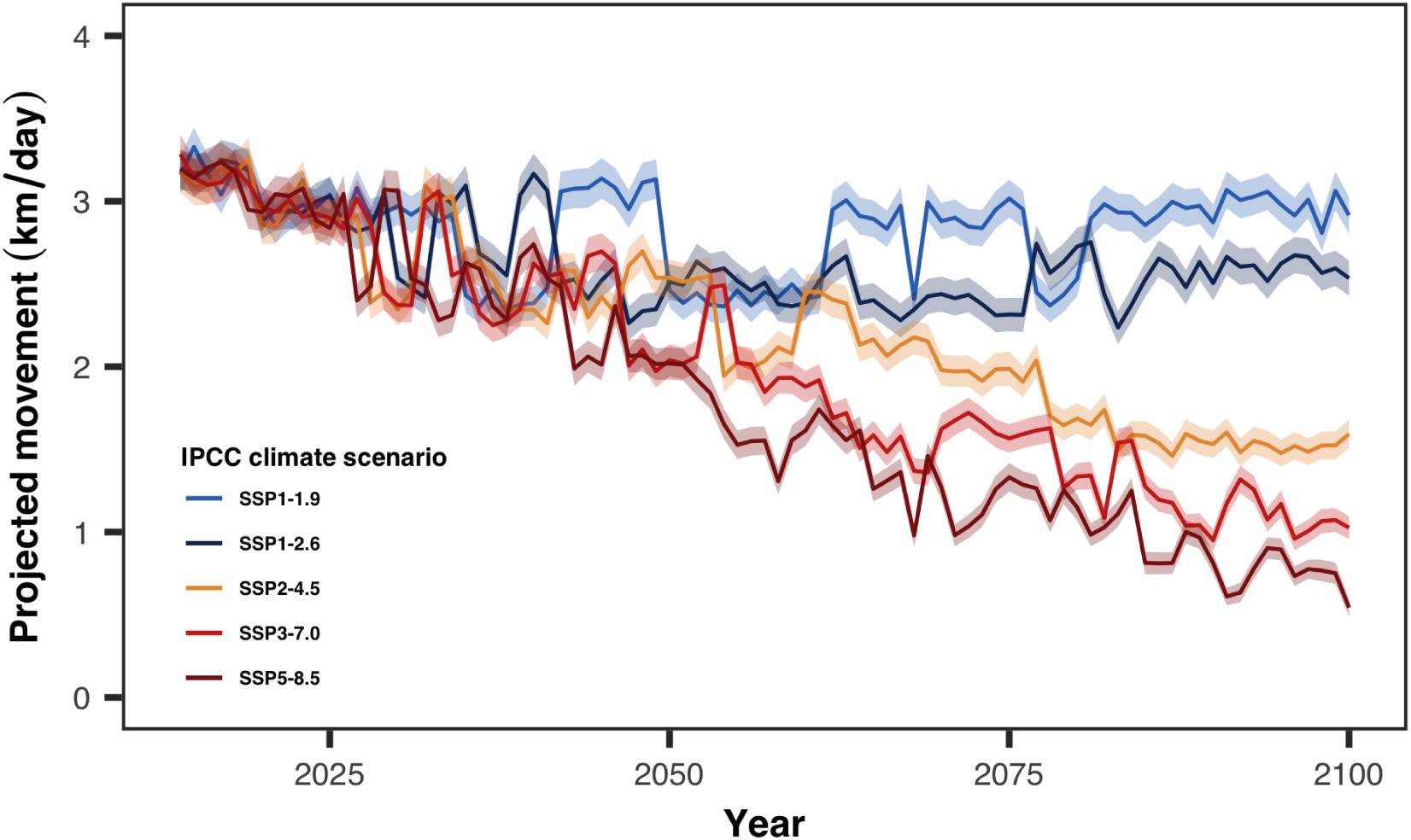
Mean annual movement of giant anteaters projected under five IPCC climate change scenarios. The solid lines depict the mean projections, whereas the shaded areas depict the 95% confidence intervals on the mean. Note how anteaters appear to be able to cope with temperatures predicted under both of the SSP1 scenarios, but show substantial reductions in movement under all of the other scenarios.

## Discussion

Giant anteater populations have suffered severe reductions in recent years and habitat loss and road-induced mortality are currently recognised as the primary threats to the species’ survival (Miranda *et al*., 2014). Our study demonstrates how giant anteater movement behaviour is tightly coupled to atmospheric temperature. As a result, the warmer temperatures predicted for Brazil throughout the rest of the 21^st^ century represent a significant threat to the species’ survival.

### Giant anteater movement and temperature

Giant anteater movement exhibited a clear quadratic response to temperature consistent with a classical ‘behavioural reaction norm’ (Dingemanse *et al*., 2010). Giant anteater movement speed peaked at 23.7°C and 95% of their movement occurred between 15.0 – 32.3°C. The lower end of this reaction norm is more or less consistent with the 17°C threshold identified by Camilo-Alves & Mourão (2006) for giant anteaters in the Pantanal. Camilo-Alves & Mourão (2006) did not identify an upper threshold for their movement and focused primarily on behavioural responses to cold stress as they monitored animals for only a 15-day period in Austral spring. The upper threshold of ∼32 °C is therefore novel to this study, and both values are roughly in line with the range of temperatures over which anteaters remain thermally neutral (∼15 – 36°C) (McNab, 1984).

While we focused on short-term weather as opposed to climate *per se*, our findings allowed us to project the behavioural responses expected under future climate change, under the assumption that giant anteaters respond to future temperatures in a way that is consistent with present-day responses. Under this assumption, the warmer temperatures predicted for Brazil throughout the rest of the 21^st^ century appear likely to challenge giant anteater survival. Importantly however, our projections indicated that while all of the SSP climate change scenarios are likely to result in decreased giant anteater movement, not all of these scenarios had the same impact. Projections for the two SSP1 scenarios suggest only modest reductions in movement that will remain relatively stable through the coming century. Projections for the higher emissions SSP2-4.5, SSP3-7.0, and SSP5-8.5 scenarios, in contrast, showed the potential for dramatic reductions in giant anteater movement, particularly in the second half of the 21^st^ century. These changes were not insubstantial. Under the SSP5-8.5 scenario we identified the potential for as much as an 84% reduction in movement speed, with giant anteaters shifting from moving an average of 3.37 km/day to only 0.55 km/day. As conditions continue to warm, individuals may experience prolonged periods of time at temperatures above their ∼32 °C thermal threshold for movement. Although giant anteaters have relatively low metabolic requirements (McNab, 1984), they cannot persist without eating indefinitely and still need to find sufficient food to grow and reproduce. Despite having a relatively low metabolic rate, giant anteaters must consume large amounts of food to sustain themselves due to the low-caloric value of their diet, which limits the durations over which they can rely on thermal refugia. Giant anteaters are a myrmecophagous specialist that consume a wide diversity of prey species associated with multiple habitats (Santana *et al*., 2024), and require extensive movement to forage. Further, giant anteaters feed for only 40 sec on each ant or termite nest on average (Redford, 1985), likely to avoid ant and termite defensive swarming responses (McNab, 1984), and must therefore constantly move to find sufficient forage. The hotter temperatures predicted to occur under the SSP2-4.5, SSP3-7.0, and SSP5-8.5 scenarios therefore represent a clear threat to the species’ persistence, as more frequent and prolonged thermally driven resting bouts are incompatible with their foraging behaviour.

### Giant anteater habitat selection and temperature

Our findings indicated that anteaters increasingly selected for native forest as temperatures increased. This partially aligns with previous studies, which have shown giant anteaters exhibit increasing selection for forests at both low and high temperatures, compared to mild ones (Camilo-Alves & Mourão, 2006, Giroux *et al*., 2023, Mourão & Medri, 2007). A possible explanation for the lack of any responses to colder weather in the present study is that cold fronts have become increasingly rare in Brazil, and our sample of temperatures below giant anteater’s thermal neutral zone (<15 °C) was very small. Despite this, our results confirm the importance of natural forest cover as thermal refugia in a warming world. This is important for a few reasons: first, it indicates that anteaters can use native forest to reduce the influence of heat on their activity. Like other species (e.g., Alston *et al*., 2020, Hovick *et al*., 2014, Sarmento *et al*., 2019), anteaters have some behavioral flexibility that allows them to use shade and other cool areas of the landscape to mitigate the negative impacts of heat stress on their physiological well-being. Accordingly, the importance of forests as a resource for giant anteaters has been shown, as these animals increase their home-range size when access to forests within their home range decreases (Giroux *et al*., 2021b). Nevertheless, anteaters still reduce their movements substantially at temperatures > 22 °C, so habitat selection is a limited solution for mitigating the negative consequences of high temperatures.

Second, eliminating native vegetation cover may reduce availability of an important habitat component for anteaters in a warmer world. Plantations of non-native eucalyptus trees have become increasingly common in the study area in recent years, and local conservationists have been concerned that this non-native forest type may not provide the same level of thermal refuge for anteaters as the tree species native to the Cerrado. Our study indicates that these fears may be well-placed—we did not find evidence that anteaters use plantations more as temperature increased (although plantations were also relatively rare on this landscape, and anteaters may visit plantations more in landscapes where they are more common). Likewise, agriculture is likely to rapidly intensify in this area in coming years, which will leave the landscape increasingly bereft of tree cover (Luiz & Steinke, 2022, Overbeck *et al*., 2022). The direct impacts on anteaters from agricultural intensification altering the supply of ants and termites is a major concern for anteaters, and the elimination of tree cover will only compound this threat.

Finally, riparian borders are legally protected from deforestation (Brasil, 2012), providing a potential refuge for anteaters even as land-use change continues into the future. Anteaters are already using these forested corridors to mitigate the negative consequences of high temperatures,but these are likely to become even more important as the native Cerrado vegetation continues to be cleared and converted to other land uses (Guidotti *et al*., 2020). Continued or even expanded protections for these stream corridors are likely to benefit anteaters as climate change progresses, and ensuring high connectivity of these important thermal buffers will be of importance for helping to ensure the species’ future.

## Conclusion

Climate change is a growing threat to wildlife. Therefore, understanding how animals respond to temperature fluctuations is critical for assessing species long-term survival and guiding conservation practice. Our study highlights the significant role that temperature plays in influencing the movement speed and habitat selection of giant anteaters. The projected increase in temperatures poses a risk of heat stress, which may drastically reduce their movement and, consequently, ability to forage, threatening their persistence. Additionally, the increased selection for native forests as temperatures rise underscores the importance of preserving these habitats, as they provide crucial thermal refuges. We highlight that giant anteaters are expected to be comparatively less sensitive to heat than other mammals of similar body mass due to their low body heat production and high thermal insulation (McNab, 1984). Therefore, it is likely that other mammals will be even more affected by rising temperatures, experiencing greater reductions in movement and an increased reliance on forests as thermal refuges (e.g., Alston *et al*., 2020, Borowik *et al*., 2020). These findings reinforce the critical role of forest conservation in mitigating the impacts of climate change on wildlife (see also De Frenne *et al*., 2019). Protecting and restoring natural habitats will be essential for ensuring the survival of diverse species as global temperatures continue to rise.

## Acknowledgements

We would like to thank the donors to the Anteaters & Highways Project especially the Foundation Segre as well as North American and European Zoos listed at (http://www.giantanteater.org/). We would also like to thank the owners of all the ranches that allowed us to monitor animals on their property, in particular Nhuveira, Quatro Irmãos and Santa Lourdes ranches. Thank you to M. Alves, D. Kluyber, C. Luba, A. Alves, and D.R. Yogui. We thank Ryan Gill for assistance in preparing figure 1. MJN was supported by an NSERC Discovery Grant RGPIN-2021-02758 and the Canadian Foundation for Innovation. AG acknowledges the Ewel postdoctoral fellowship support and the Brazilian National Council for Scientific and Technological Development - CNPq (317355/2023-6)

## Data Accessibility

The GPS tracking data used in this analysis are openly available on the Movebank data repository (Movebank ID: 1574830796). The climate change projections are openly available via World Bank’s Climate Knowledge Portal. The R scripts used to carry out this study are openly available on GitHub at https://github.com/integratecology/anteaters.

## Authors’ Contributions

MJN, JMA, and ALJD conceived the ideas; JMA and MJN conducted the analyses; and JMA, MJN, AG, and ALJD led the writing of the manuscript. All authors contributed critically to drafts of the manuscript and gave final approval for publication.

## Supplementary materials

**Figure S1:**
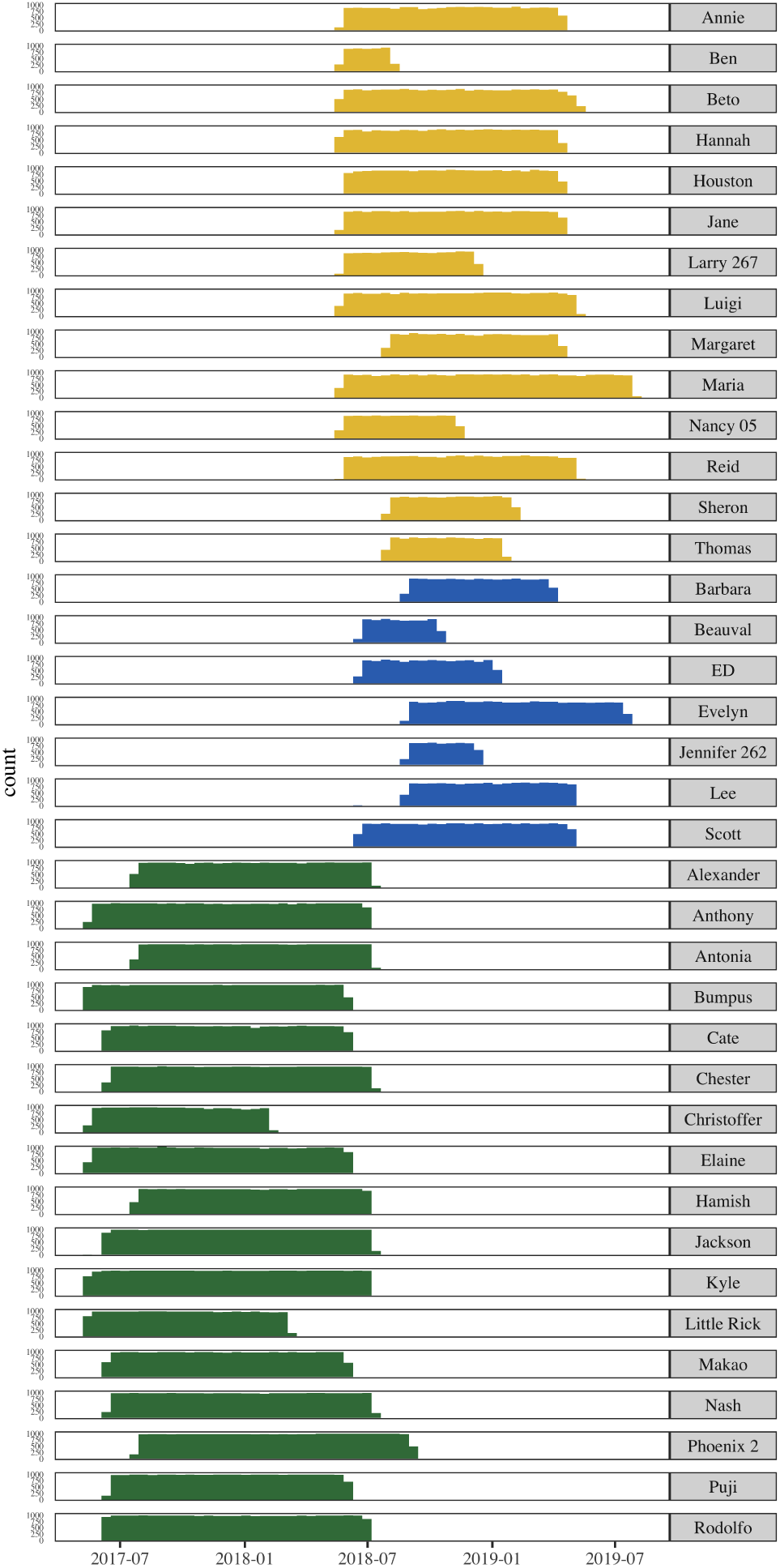
Histograms depicting of the amount of data collected over time for each of the 38 individuals included in the analyses presented in the main text. The different colours correspond to the three different study sites.

**Figure S2:**
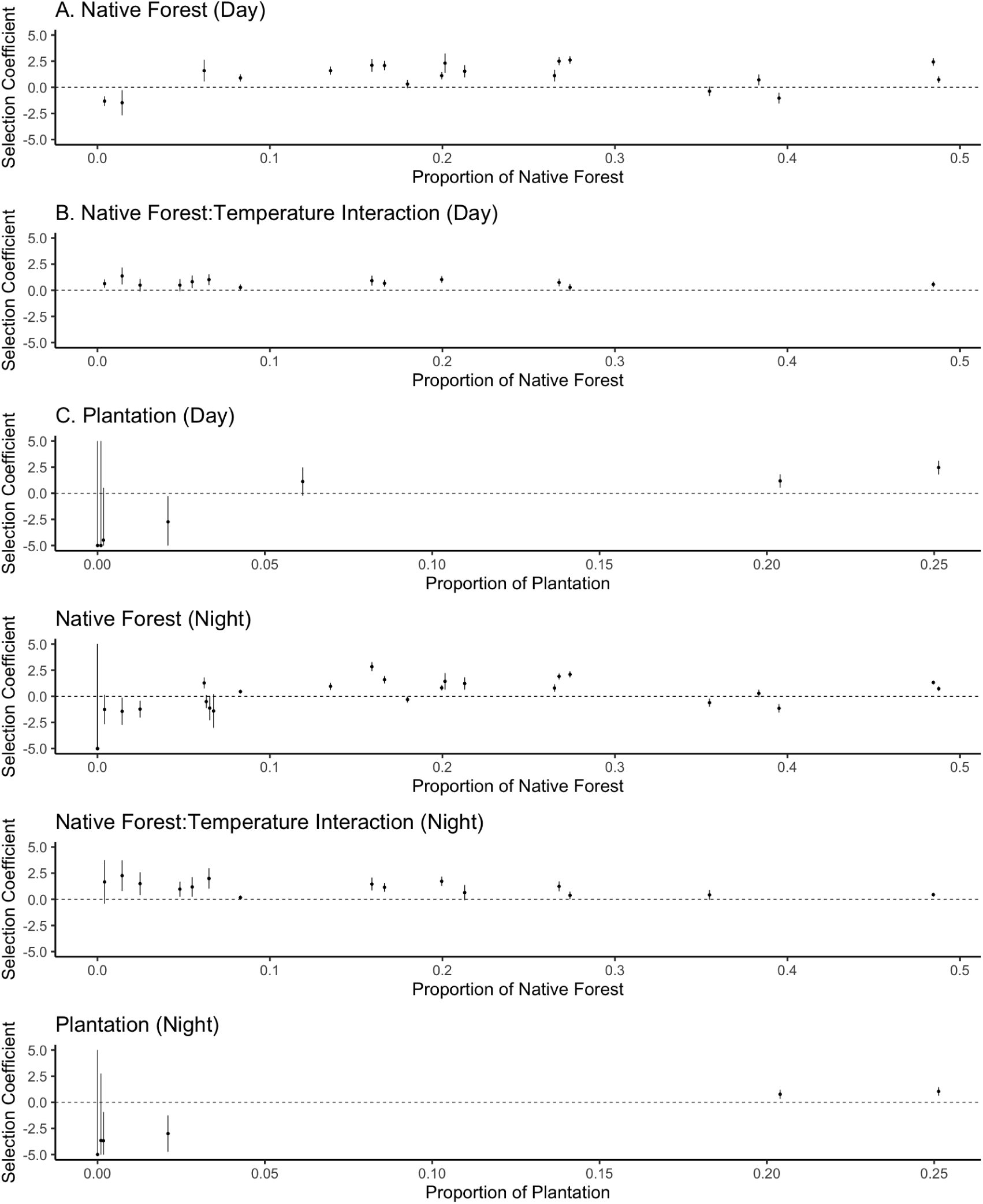
Graphs depicting parameter estimates in individual resource selection functions at varying amounts of visitation of native forest and plantation for each of the 38 individuals included in the analyses presented in the main text. Bars represent 95% confidence intervals. There are fewer than 38 points in each plot because individual parameters were often missing from the most-supported model for each individual.

## Notes

https://github.com/integratecology/anteaters

### Competing Interest Statement

The authors have declared no competing interest.

## References

Alston, J.M., Dillon, M.E., Keinath, D.A., Abernethy, I.M. & Goheen, J.R. (2022) Daily torpor reduces the energetic consequences of microhabitat selection for a widespread bat. Ecology 103, e3677.

Alston, J.M., Fleming, C.H., Kays, R., Streicher, J.P., Downs, C.T., Ramesh, T., Reineking, B. & Calabrese, J.M. (2023) Mitigating pseudoreplication and bias in resource selection functions with autocorrelation-informed weighting. Methods in Ecology and Evolution 14, 643–654.

Alston, J.M., Joyce, M.J., Merkle, J.A. & Moen, R.A. (2020) Temperature shapes movement and habitat selection by a heat-sensitive ungulate. Landscape Ecology 35, 1961–1973.

Alvares, C.A., Stape, J.L., Sentelhas, P.C., Gonçalves, J.d.M., Sparovek, G., et al. (2013) Köppen’s climate classification map for brazil. Meteorologische zeitschrift 22, 711–728.

Avgar, T., Potts, J.R., Lewis, M.A. & Boyce, M.S. (2016) Integrated step selection analysis: bridging the gap between resource selection and animal movement. Methods in Ecology and Evolution 7, 619–630.

Barnosky, A.D., Matzke, N., Tomiya, S., Wogan, G.O.U., Swartz, B., Quental, T.B., Marshall, C., McGuire, J.L., Lindsey, E.L., Maguire, K.C., Mersey, B. & Ferrer, E.A. (2011) Has the Earth’s sixth mass extinction already arrived? Nature 471, 51–57.

Beever, E.A., Hall, L.E., Varner, J., Loosen, A.E., Dunham, J.B., Gahl, M.K., Smith, F.A. & Lawler, J.J. (2017) Behavioral flexibility as a mechanism for coping with climate change. Frontiers in Ecology and the Environment 15, 299–308.

Borowik, T., Ratkiewicz, M., Maślanko, W., Duda, N. & Kowalczyk, R. (2020) Too hot to handle: summer space use shift in a cold-adapted ungulate at the edge of its range. Landscape Ecology 35, 1341–1351.

Brasil, N.C.F. (2012) Lei n 12.651, de 25 de maio de 2012. Brasília, Diário Oficial da União http://www.planalto.gov.br/ccivil_03/_ato2011-2014/2012/lei/l12651.htm.

Buchholz, R., Banusiewicz, J.D., Burgess, S., Crocker-Buta, S., Eveland, L. & Fuller, L. (2019) Behavioural research priorities for the study of animal response to climate change. Animal Behaviour 150, 127–137.

Calabrese, J.M., Fleming, C.H. & Gurarie, E. (2016) ctmm: an r package for analyzing animal relocation data as a continuous-time stochastic process. Methods in Ecology and Evolution 7, 1124–1132.

Camilo-Alves, C.d.S.e.P. & Mourão, G.d.M. (2006) Responses of a specialized insectivorous mammal (*Myrmecophaga tridactyla*) to variation in ambient temperature. Biotropica 38, 52–56.

Ceballos, G., Ehrlich, P.R., Barnosky, A.D., García, A., Pringle, R.M. & Palmer, T.M. (2015) Accelerated modern human–induced species losses: Entering the sixth mass extinction. Science Advances 1, e1400253.

Ceballos, G., Ehrlich, P.R. & Dirzo, R. (2017) Biological annihilation via the ongoing sixth mass extinction signaled by vertebrate population losses and declines. Proceedings of the National Academy of Sciences 114, E6089–E6096.

Cunningham, G.D., While, G.M., Olsson, M., Ljungström, G. & Wapstra, E. (2020) Degrees of change: between and within population variation in thermal reaction norms of phenology in a viviparous lizard. Ecology 101, e03136.

De Frenne, P., Zellweger, F., Rodríguez-Sánchez, F., Scheffers, B.R., Hylander, K., Luoto, M., Vellend, M., Verheyen, K. & Lenoir, J. (2019) Global buffering of temperatures under forest canopies. Nature Ecology & Evolution 3, 744–749.

Dingemanse, N.J., Kazem, A.J.N., Réale, D. & Wright, J. (2010) Behavioural reaction norms: animal personality meets individual plasticity. Trends in Ecology & Evolution 25, 81–89.

Fleming, C.H., Calabrese, J.M., Mueller, T., Olson, K.A., Leimgruber, P. & Fagan, W.F. (2014) From fine-scale foraging to home ranges: a semivariance approach to identifying movement modes across spatiotemporal scales. American Naturalist 183, E154–E167.

Fleming, C.H., Drescher-Lehman, J., Noonan, M.J., Akre, T.S.B., Brown, D.J., Cochrane, M.M., Dejid, N., DeNicola, V., DePerno, C.S., Dunlop, J.N., Gould, N.P., Harrison, A.L., Hollins, J., Ishii, H., Kaneko, Y., Kays, R., Killen, S.S., Koeck, B., Lambertucci, S.A., LaPoint, S.D., Medici, E.P., Meyburg, B.U., Miller, T.A., Moen, R.A., Mueller, T., Pfeiffer, T., Pike, K.N., Roulin, A., Safi, K., Séchaud, R., Scharf, A.K., Shephard, J.M., Stabach, J.A., Stein, K., Tonra, C.M., Yamazaki, K., Fagan, W.F. & Calabrese, J.M. (2021) A comprehensive framework for handling location error in animal tracking data.

Fleming, C.H., Noonan, M.J., Medici, E.P. & Calabrese, J.M. (2019) Overcoming the challenge of small effective sample sizes in home-range estimation. Methods in Ecology and Evolution 10, 1679–1689.

Fleming, C.H., Sheldon, D., Fagan, W.F., Leimgruber, P., Mueller, T., Nandintsetseg, D., Noonan, M.J., Olson, K.A., Setyawan, E., Sianipar, A. & Calabrese, J.M. (2018) Correcting for missing and irregular data in home-range estimation. Ecological Applications 28, 1003–1010.

Gaudin, T.J., Hicks, P. & Di Blanco, Y. (2018) *Myrmecophaga tridactyla* (Pilosa: Myrmecophagidae). Mammalian Species 50, 1–13.

Giroux, A., Ortega, Z., Attias, N., Desbiez, A.L.J., Valle, D., Börger, L. & Oliveira-Santos, L.G.R. (2023) Activity modulation and selection for forests help giant anteaters to cope with temperature changes. Animal Behaviour 201, 191–209.

Giroux, A., Ortega, Z., Bertassoni, A., Desbiez, A.L.J., Kluyber, D., Massocato, G.F., Miranda, G.D., Mourão, G., Surita, L., Attias, N., Bianchi, R.d.C., Gasparotto, V.P.d.O. & Oliveira-Santos, L.G.R. (2021a) The role of environmental temperature on movement patterns of giant anteaters. Integrative Zoology **n/a**.

Giroux, A., Ortega, Z., Oliveira-Santos, L.G.R., Attias, N., Bertassoni, A. & Desbiez, A.L.J. (2021b) Sexual, allometric and forest cover effects on giant anteaters’ movement ecology. PLoS One 16, e0253345.

Guidotti, V., de Barros Ferraz, S.F., Pinto, L.F.G., Sparovek, G., Taniwaki, R.H., Garcia, L.G. & Brancalion, P.H. (2020) Changes in brazil’s forest code can erode the potential of riparian buffers to supply watershed services. Land Use Policy 94, 104511.

Haase, C.G., Fletcher Jr, R.J., Slone, D.H., Reid, J.P. & Butler, S.M. (2020) Traveling to thermal refuges during stressful temperatures leads to foraging constraints in a central-place forager. Journal of Mammalogy 101, 271–280.

Hovick, T.J., Elmore, R.D., Allred, B.W., Fuhlendorf, S.D. & Dahlgren, D.K. (2014) Landscapes as a moderator of thermal extremes: a case study from an imperiled grouse. Ecosphere 5, 1–12.

Huey, R.B., Kearney, M.R., Krockenberger, A., Holtum, J.A.M., Jess, M. & Williams, S.E. (2012) Predicting organismal vulnerability to climate warming: roles of behaviour, physiology and adaptation. Philosophical Transactions of the Royal Society B: Biological Sciences 367, 1665–1679.

Johnson, D.H. (1980) The comparison of usage and availability measurements for evaluating resource preference. Ecology 61, 65–71.

Kearney, M., Shine, R. & Porter, W.P. (2009) The potential for behavioral thermoregulation to buffer “cold-blooded” animals against climate warming. Proceedings of the National Academy of Sciences 106, 3835–3840.

Kleiber, M. (1932) Body size and metabolism. Hilgardia 6, 315–353.

Kluyber, D., Attias, N., Alves, M.H., Alves, A.C., Massocato, G. & Desbiez, A.L.J. (2021) Physical capture and chemical immobilization procedures for a mammal with singular anatomy: the giant anteater (*Myrmecophaga tridactyla*). European Journal of Wildlife Research 67, 67.

Levy, O., Dayan, T., Porter, W.P. & Kronfeld-Schor, N. (2019) Time and ecological resilience: can diurnal animals compensate for climate change by shifting to nocturnal activity? Ecological Monographs 89, e01334.

Long, R.A., Bowyer, R.T., Porter, W.P., Mathewson, P., Monteith, K.L. & Kie, J.G. (2014) Behavior and nutritional condition buffer a large-bodied endotherm against direct and indirect effects of climate. Ecological Monographs 84, 513–532.

Luiz, C.H.P. & Steinke, V.A. (2022) Recent environmental legislation in brazil and the impact on cerrado deforestation rates. Sustainability 14, 8096.

Mantyka-Pringle, C.S., Visconti, P., Di Marco, M., Martin, T.G., Rondinini, C. & Rhodes, J.R. (2015) Climate change modifies risk of global biodiversity loss due to land-cover change. Biological Conservation 187, 103–111.

Masson-Delmotte, V., Zhai, P., Pirani, A., Connors, S.L., Péan, C., Berger, S., Caud, N., Chen, Y., Goldfarb, L., Gomis, M.I., Huang, M., Leitzell, K., Lonnoy, E., Matthews, J.B.R., Maycock, T.K., Waterfield, T., Yelekçi Yu, R. & Zhou, B. (eds.) (2021) Climate Change 2021: The Physical Science Basis. Contribution of Working Group I to the Sixth Assessment Report of the Intergovernmental Panel on Climate Change. Cambridge University Press, New York, NY, USA.

McNab, B.K. (1984) Physiological convergence amongst ant-eating and termite-eating mammals. Journal of Zoology 203, 485–510.

McNab, B.K. (1986) The influence of food habits on the energetics of eutherian mammals. Ecological Monographs 56, 1–19.

Miranda, F., Bertassoni, A. & Abba, A.M. (2014) Myrmecophaga tridactyla. Tech. Rep. e.T14224A47441961.

Moritz, C. & Agudo, R. (2013) The future of species under climate change: resilience or decline? Science 341, 504–508.

Mourão, G. & Medri, Í. (2007) Activity of a specialized insectivorous mammal (myrmecophaga tridactyla) in the pantanal of brazil. Journal of zoology 271, 187–192.

Muñoz-Sabater, J., Dutra, E., Agustí-Panareda, A., Albergel, C., Arduini, G., Balsamo, G., Boussetta, S., Choulga, M., Harrigan, S., Hersbach, H. et al. (2021) Era5-land: A state-of-the-art global reanalysis dataset for land applications. Earth system science data 13, 4349–4383.

Noonan, M.J., Ascensão, F., Yogui, D.R. & Desbiez, A.L.J. (2022) Roads as ecological traps for giant anteaters. Animal Conservation 25, 182–194.

Noonan, M.J., Fleming, C.H., Akre, T.S., Drescher-Lehman, J., Gurarie, E., Harrison, A.L., Kays, R. & Calabrese, J.M. (2019a) Scale-insensitive estimation of speed and distance traveled from animal tracking data. Movement Ecology 7, 35.

Noonan, M.J., Markham, A., Newman, C., Trigoni, N., Buesching, C.D., Ellwood, S.A. & Macdonald, D.W. (2014) Climate and the individual: inter-annual variation in the autumnal activity of the European badger (*Meles meles*). PLOS ONE 9, e83156, publisher: Public Library of Science.

Noonan, M.J., Newman, C., Markham, A., Bilham, K., Buesching, C.D. & Macdonald, D.W. (2018) In situ behavioral plasticity as compensation for weather variability: implications for future climate change. Climatic Change 149, 457–471.

Noonan, M.J., Tucker, M.A., Fleming, C.H., Akre, T.S., Alberts, S.C., Ali, A.H., Altmann, J., Antunes, P.C., Belant, J.L., Beyer, D., Blaum, N., Böhning-Gaese, K., Cullen, L., Paula, R.C.d., Dekker, J., Drescher-Lehman, J., Farwig, N., Fichtel, C., Fischer, C., Ford, A.T., Goheen, J.R., Janssen, R., Jeltsch, F., Kauffman, M., Kappeler, P.M., Koch, F., LaPoint, S., Markham, A.C., Medici, E.P., Morato, R.G., Nathan, R., Oliveira-Santos, L.G.R., Olson, K.A., Patterson, B.D., Paviolo, A., Ramalho, E.E., Rösner, S., Schabo, D.G., Selva, N., Sergiel, A., Silva, M.X.d., Spiegel, O., Thompson, P., Ullmann, W., Zięba, F., Zwijacz-Kozica, T., Fagan, W.F., Mueller, T. & Calabrese, J.M. (2019b) A comprehensive analysis of autocorrelation and bias in home range estimation. Ecological Monographs 89, e01344.

O’Leary, M.A., Bloch, J.I., Flynn, J.J., Gaudin, T.J., Giallombardo, A., Giannini, N.P., Goldberg, S.L., Kraatz, B.P., Luo, Z.X., Meng, J., Ni, X., Novacek, M.J., Perini, F.A., Randall, Z.S., Rougier, G.W., Sargis, E.J., Silcox, M.T., Simmons, N.B., Spaulding, M., Velazco, P.M., Weksler, M., Wible, J.R. & Cirranello, A.L. (2013) The placental mammal ancestor and the post–K-Pg radiation of placentals. Science 339, 662–667.

Overbeck, G.E., Vélez-Martin, E., da Silva Menezes, L., Anand, M., Baeza, S., Carlucci, M.B., Dechoum, M.S., Durigan, G., Fidelis, A., Guido, A. et al. (2022) Placing brazil’s grasslands and savannas on the map of science and conservation. *Perspectives in Plant Ecology*, Evolution and Systematics 56, 125687.

Prothero, D.R. (1994) The late Eocene-Oligocene extinctions. Annual Review of Earth and Planetary Sciences 22, 145–165.

R Core Team, (2020) R: a language and environment for statistical computing.

Redford, K.H. (1985) Feeding and food preference in captive and wild giant anteaters (*Myrmecophaga tridactyla*). Journal of Zoology 205, 559–572.

Riahi, K., van Vuuren, D.P., Kriegler, E., Edmonds, J., O’Neill, B.C., Fujimori, S., Bauer, N., Calvin, K., Dellink, R., Fricko, O., Lutz, W., Popp, A., Cuaresma, J.C., Kc, S., Leimbach, M., Jiang, L., Kram, T., Rao, S., Emmerling, J., Ebi, K., Hasegawa, T., Havlik, P., Humpenöder, F., Da Silva, L.A., Smith, S., Stehfest, E., Bosetti, V., Eom, J., Gernaat, D., Masui, T., Rogelj, J., Strefler, J., Drouet, L., Krey, V., Luderer, G., Harmsen, M., Takahashi, K., Baumstark, L., Doelman, J.C., Kainuma, M., Klimont, Z., Marangoni, G., Lotze-Campen, H., Obersteiner, M., Tabeau, A. & Tavoni, M. (2017) The Shared Socioeconomic Pathways and their energy, land use, and greenhouse gas emissions implications: an overview. Global Environmental Change 42, 153–168.

Santana, T.G., Attias, N., Nascimento, N.T., Tibcherani, M., Rocha, M.M. & Desbiez, A.L.J. (2024) No evidence of sex-related differences in the diet of giant anteater in the brazilian savanna. Mammalian Biology pp. 1–12.

Sarmento, W., Biel, M. & Berger, J. (2019) Seeking snow and breathing hard – behavioral tactics in high elevation mammals to combat warming temperatures. PLOS ONE 14.

Sheppard, A., Hecker, L., Edwards, M. & Nielsen, S. (2021) Determining the influence of snow and temperature on the movement rates of wood bison (*Bison bison athabascae*). Canadian Journal of Zoology 99, 489–496.

Sih, A., Ferrari, M.C.O. & Harris, D.J. (2011) Evolution and behavioural responses to human-induced rapid environmental change. Evolutionary Applications 4, 367–387.

Souza, C.M., Z. Shimbo, J., Rosa, M.R., Parente, L.L. A. Alencar, A., Rudorff, B.F.T., Hasenack, H., Matsumoto, M. G. Ferreira, L., Souza-Filho, P.W.M., de Oliveira, S.W., Rocha, W.F., Fonseca, A.V., Marques, C.B., Diniz, C.G., Costa, D., Monteiro, D., Rosa, E.R., Vélez-Martin, E., Weber, E.J., Lenti, F.E.B., Paternost, F.F., Pareyn, F.G.C., Siqueira, J.V., Viera, J.L., Neto, L.C.F., Saraiva, M.M., Sales, M.H., Salgado, M.P.G., Vasconcelos, R., Galano, S., Mesquita, V.V. & Azevedo, T. (2020) Reconstructing three decades of land use and land cover changes in Brazilian biomes with Landsat Archive and Earth Engine. Remote Sensing 12, 2735.

Sunday, J.M., Bates, A.E., Kearney, M.R., Colwell, R.K., Dulvy, N.K., Longino, J.T. & Huey, R.B. (2014) Thermal-safety margins and the necessity of thermoregulatory behavior across latitude and elevation. Proceedings of the National Academy of Sciences p. 201316145.

Urban, M.C. (2015) Accelerating extinction risk from climate change. Science 348, 571–573.

Venter, O., Brodeur, N.N., Nemiroff, L., Belland, B., Dolinsek, I.J. & Grant, J.W.A. (2006) Threats to endangered species in Canada. BioScience 56, 903–910.

Wiens, J.J. (2016) Climate-related local extinctions are already widespread among plant and animal species. PLOS Biology 14, e2001104.

Woo-Durand, C., Matte, J.M., Cuddihy, G., McGourdji, C.L., Venter, O. & Grant, J.W.A. (2020) Increasing importance of climate change and other threats to at-risk species in Canada. Environmental Reviews.

